# Checkpoint signaling and error correction require regulation of the MPS1 T-loop by PP2A-B56

**DOI:** 10.1101/626937

**Authors:** Daniel Hayward, James Bancroft, Davinderpreet Mangat, Tatiana Alfonso-Pérez, Sholto Dugdale, Julia McCarthy, Francis A. Barr, Ulrike Gruneberg

## Abstract

During mitosis, the formation of microtubule-kinetochore attachments is monitored by the serine/threonine kinase Mono-Polar Spindle 1 (MPS1). MPS1 is recruited to unattached kinetochores where it phosphorylates KNL1, BUB1 and MAD1 to initiate the spindle checkpoint. This arrests the cell cycle until all kinetochores have been stably captured by microtubules. MPS1 also contributes to the error correction process rectifying incorrect kinetochore attachments. MPS1 activity at kinetochores requires auto-phosphorylation at multiple sites including T676 in the activation segment or “T-loop”. We now demonstrate that a BUBR1-bound pool of PP2A-B56 regulates MPS1 T-loop autophosphorylation and hence activation status in mammalian cells. Overriding this regulation using phospho-mimetic mutations in the MPS1 T-loop to generate a constitutively active kinase results in a prolonged mitotic arrest with continuous turn-over of microtubule-kinetochore attachments. Dynamic regulation of MPS1 catalytic activity by kinetochore-localized PP2A-B56 is thus critical for controlled MPS1 activity and timely cell cycle progression.

## Introduction

Unattached kinetochores are detected by the spindle assembly checkpoint (SAC) (Musacchio, 2015). During mitosis, generation of a checkpoint response, and recruitment of all other SAC proteins to kinetochores depends on the activity of a conserved protein kinase, MPS1 (Ciliberto and Hauf, 2017; Liu and Winey, 2012; Musacchio, 2015; Pachis and Kops, 2018). MPS1 is phosphorylated by CDK1-CCNB1 at S281 and then localises to kinetochores as soon as the nuclear envelope breaks down and is lost from kinetochores as they attach (Hayward et al., 2019; Jelluma et al., 2010; Stucke et al., 2002). MPS1 binds to kinetochores via multiple weak interactions with the HEC1/NDC80 complex in a manner competitive with microtubule binding (Hiruma et al., 2015; Ji et al., 2015). This supports a simple model explaining the restriction of MPS1 activity to mitosis by the requirement for CDK1-CCNB1 and how MPS1-dependent checkpoint signaling is coupled to microtubule binding.

MPS1 activation at kinetochores requires autophosphorylation of the T-loop on threonine 676 (T676) (Jelluma et al., 2008a; Kang et al., 2007; Mattison et al., 2007). Like other kinases, MPS1 is thought to trans auto-phosphorylate and thus self-activate, an event that would be promoted by clustering of several MPS1 molecules at an unattached kinetochore (Combes et al., 2018; Dodson et al., 2013; Kang et al., 2007). Kinetochore localisation and activity of MPS1 are therefore linked. Inhibition of MPS1 activity by small molecule inhibitors or mutations in the MPS1 active site result in increased rather than decreased levels of MPS1 at kinetochores (Hewitt et al., 2010; Jelluma et al., 2010; Santaguida et al., 2010). This is not simply explained. However, autophosphorylation of the N-terminus of MPS1, in addition to relieving autoinhibition of kinase activity, has been suggested to promote release from the kinetochore, possibly by interfering with Hec1 binding (Combes et al., 2018; Wang et al., 2014).

Since initiation of the spindle assembly checkpoint requires multi-site phosphorylation of KNL1, MAD1 and BUB1 by MPS1 as well as MPS1 phosphorylation by multiple kinases, termination of the checkpoint response must require phosphatases. Both PP2A-B56 and PP1 have been implicated in the process of dephosphorylating KNL1 and promoting spindle assembly checkpoint silencing (Espert et al., 2014; Nijenhuis et al., 2014). However, the phosphatases acting on MPS1 and other checkpoint proteins still need to be clarified. PP1 has been implicated in dephosphorylating the MPS1 T-loop in flies (Moura et al., 2017), although it is not clear whether this mechanism is conserved in mammals. PP2A-B56 exists in several spatially distinct populations in mitotic cells (Qian et al., 2013; Vallardi et al., 2019). One pool is bound to the C-terminal domain of the BUBR1 protein via a conserved LxxIxE motif (Kruse et al., 2013; Suijkerbuijk et al., 2012; Xu et al., 2013). This pool of PP2A-B56 has been shown to oppose both the Aurora B and the MPS1 kinases in chromosome alignment and spindle checkpoint signaling, respectively. Depletion of PP2A-B56 therefore results in mitotically arrested cells with unattached kinetochores (Espert et al., 2014; Foley et al., 2011; Maciejowski et al., 2017). In addition to orchestrating spindle checkpoint signaling, MPS1 also contributes directly to the turnover of erroneous microtubule-kinetochore attachments by phosphorylating the Ska complex at microtubule-kinetochore junctions. Again, this activity of MPS1 is opposed by PP2A-B56 (Maciejowski et al., 2017).

Here we demonstrate that a BUBR1-dependent pool of PP2A-B56 is a key MPS1 T-loop phosphatase. Furthermore, we investigate the role of dynamic turnover of MPS1 T-loop phosphorylation by PP2A-B56 in checkpoint signaling and error correction.

## Results and Discussion

### MPS1 T-loop phosphorylation is controlled by PP2A

MPS1 activity is dynamically regulated by autophosphorylation at Threonine 676 (T676) in the T-loop of the kinase domain (Jelluma et al., 2008a; Kang et al., 2007; Mattison et al., 2007). To identify the class of phosphatase acting at this site, mitotic HeLa cells expressing endogenously tagged MPS1-GFP were pre-treated with PPP family phosphatase inhibitors, then briefly incubated with MPS1 inhibitor (MPS1i) to stop T-loop autophosphorylation (Choy et al., 2017; Hewitt et al., 2010; Ishihara et al., 1989; Mitsuhashi et al., 2001). In control cells, MPS1i addition resulted in the loss of the MPS1 pT676 signal (Figure 1A and 1B, Figure S1A and S1B). The signal for total MPS1-GFP increased, in line with reports by other groups (Hewitt et al., 2010; Jelluma et al., 2010). Addition of the dual PP1/2A inhibitor (PP1/2Ai; calyculin) but not PP1 inhibitor (PP1i; tautomycetin), prevented the loss of the T-loop phosphorylation (Figure 1A and 1B). Neither treatment affected the increase of MPS1-GFP levels at kinetochores upon MPS1 inhibition (Figure 1A and 1C). These observations suggest that in mammalian cells, in contrast to Drosophila, a PPP-family phosphatase other than PP1 was involved in the turnover of the MPS1 T-loop phosphorylation.

**Figure 1.**
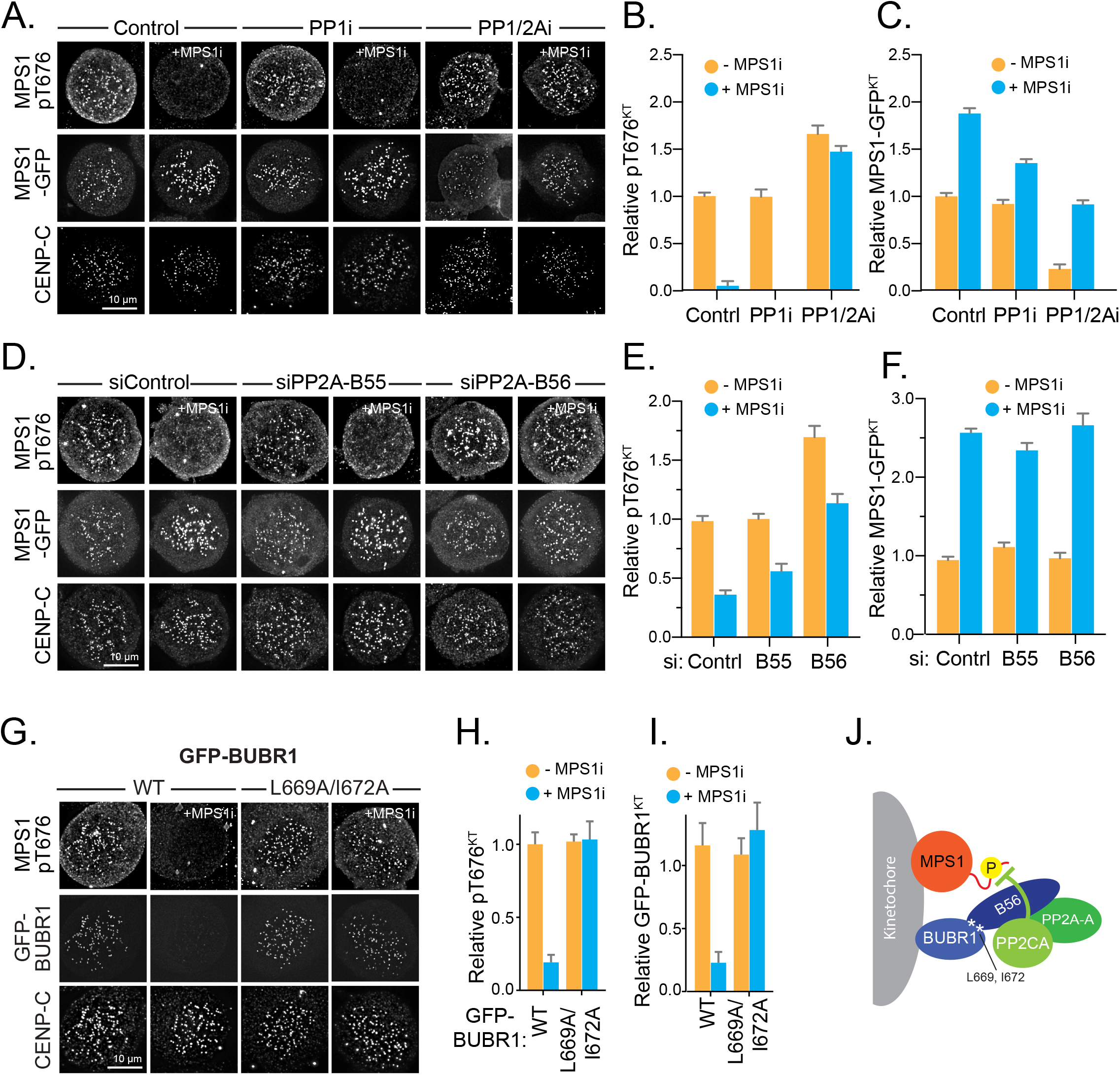
BUBR1-bound PP2A-B56 dephosphorylates MPS1 auto-activatory site T676. **(A)** Mitotic HeLa MPS1-GFP cells were pre-treated with DMSO (Control), 5 μM Tautomycetin (PP1i) or 25 nM Calyculin (PP1/PP2Ai), then 2μM MPS1 inhibitor was added 5 min prior to fixation where indicated (+MPS1i). MPS1 pT676 and CENP-C were detected using antibodies, and total MPS1 was detected using GFP fluorescence. **(B)** Mean kinetochores levels ± SEM of MPS1 pT676 phosphorylation and **(C)** MPS1 relative to the - MPS1i control are plotted. **(D)** MPS1 pT676 phosphorylation and MPS1-GFP localisation in control, PP2A-B55 or PP2A-B56 depleted mitotically arrested HeLa MPS1-GFP cells treated with MPS1i. MPS1 pT676 and CENP-C were detected using antibodies, MPS1 by GFP fluorescence. **(E)** Relative levels ± SEM of kinetochore associated MPS1 pT676 and **(F)** total MPS1-GFP are plotted. **(G)** HeLa-Flp-In/TREx GFP-BUBR1^WT^ or GFP-BUBR1^L669A/I672A^ cells were depleted of endogenous BUBR1 and transgenes were induced. Cells were mitotically arrested and DMSO (Control) or MPS1i added 5 min prior to fixation. MPS1 pT676 and CENP-C were detected using antibodies, BubR1 was detected using GFP fluorescence. **(H)** MPS1 pT676 phosphorylation and **(I)** Mean GFP-BUBR1 kinetochore levels ± SEM are plotted. >10 kinetochores were measured per cell in >15 cells from two independent experiments. **(J)** A model illustrating how BUBR1-associated PP2A-B56 regulates the MPS1 T-loop auto-phosphorylation.

To identify the PPP family phosphatase responsible for the turnover of the pT676 site, GFP-MPS1 cells were depleted of individual PPP family catalytic subunits using siRNA (Espert et al., 2014). Depletion of the PP2A catalytic a subunit, the predominant PP2A catalytic subunit in HeLa cells (Janssens and Goris, 2001), resulted in a significant retention of the pT676 signal following MPS1 inhibition, whereas none of the other depletions, including PP1α, β and γ co-depletions, had the same effect (Figure S1C-S1F). This result was independent of whether cells had been arrested in mitosis or were passing through an unperturbed cell cycle (Figure 2SA and 2SB). These data confirm the results of the phosphatase inhibitor treatments (Figure 1A-1C) and suggest that a PP2A phosphatase holoenzyme complex is responsible for modulating the MPS1-T676 phosphorylation.

**Figure 2.**
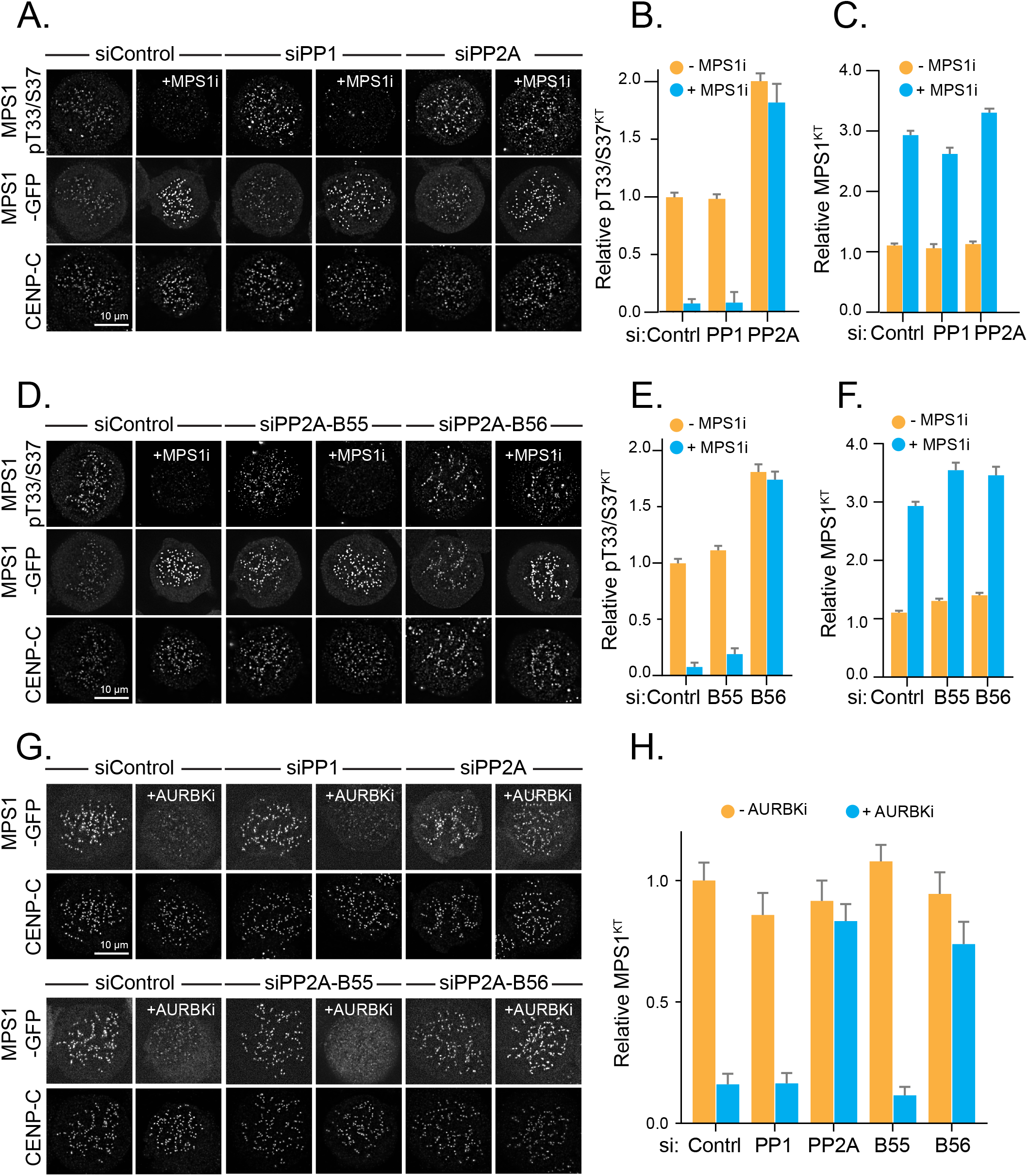
PP2A-B56 modulates MPS1 kinetochore localization by opposing both MPS1 and Aurora B. **(A)** MPS1 pT33/S37 phosphorylation, MPS1 and CENP-C localisation were compared before and after MPS1i in control, PP1αβγ or PP2A catalytic α subunit depleted HeLa MPS1-GFP cells. **(B)** Mean kinetochore levels ±SEM of pT33/S37 and **(C)** total MPS1-GFP relative to CENP-C are plotted. **(D)** MPS1 pT33/S37 phosphorylation and MPS1 localisation were detected in control, PP2A-B55 or PP2A-B56 depleted cells. **(E)** Mean pT33/S37 and **(F)** total MPS1-GFP kinetochore levels ± SEM relative to CENP-C are plotted. **(G)** MPS1 and CENP-C localisation before and after 10 min AURBKi in Control, PP1, PP2A catalytic subunit *a* or PP2A-B55 or PP2A-B56 regulatory subunit depleted HeLa MPS1-GFP cells. **(H)** Mean kinetochore levels ± SEM of MPS1-GFP relative to CENP-C were plotted. >10 kinetochores were measured per cell in >15 cells from two independent experiments.

### BUBR1-bound PP2A-B56 regulates MPS1 T676 phosphorylation

Since the BUBR1-associated pool of the PP2A-B56 phosphatase had already been identified as the phosphatase acting on some MPS1 autophosphorylations and on MPS1 substrates at the kinetochore (Espert et al., 2014; Maciejowski et al., 2017; Qian et al., 2017), this was the most likely candidate for the MPS1 T676 phosphatase. Indeed, depletion of all B56 subunits resulted in retention of the pT676 staining upon MPS1 inhibition whereas depletion of all B55 subunits had no effect (Figure 1D and 1E, Figure S1H). The increase in kinetochore-bound MPS1 was not altered by depletion of either PP2A phosphatase (Figure 1D and 1F). To define the required pool of PP2A-B56, BUBR1 was replaced with a mutant carrying two mutations in the PP2A-B56 binding site (BUBR1^L669A/I672A^) (Espert et al., 2014; Kruse et al., 2013). In cells expressing GFP-BUBR1^L669A/I672A^, but not wild type BUBR1, MPS1 T-loop phosphorylation was retained upon MPS1i treatment (Figure 1G-1I and Figure S1I). Together, these data show that during mitosis the MPS1 T-loop phosphorylation is dynamically controlled by the BUBR1-bound pool of PP2A-B56 (Figure 1J).

### PP2A-B56 modulates MPS1 kinetochore localization by counteracting both MPS1 and Aurora B

MPS1 localization to unattached kinetochores is promoted by the CDK1-CCNB1 and Aurora B kinases and is opposed by its own activity (Hayward et al., 2019; Hewitt et al., 2010; Jelluma et al., 2010; Nijenhuis et al., 2013; Saurin et al., 2011; Zhu et al., 2013). It has been suggested that release of active MPS1 from kinetochores is mediated by auto-phosphorylation of the MPS1 N-terminus (Wang et al., 2014). We tested whether the phosphorylation status of T33 and S37 in the N-terminus of MPS1 was also regulated by PP2A-B56. In agreement with a published report, depletion of PP2A-B56 stabilized pT33/pS37 in the absence of MPS1 activity (Figure 2A-2F) (Maciejowski et al., 2017). Surprisingly, this did not result in decreased total MPS1-GFP at kinetochores (Figure 2D-2F). Aurora B is a major positive regulator of MPS1 localization to kinetochores (Nijenhuis et al., 2013; Saurin et al., 2011; Zhu et al., 2013), suggesting that PP2A-B56 may be simultaneously counteracting the MPS1 N-terminal autophosphorylations promoting MPS1 kinetochore release and the Aurora B-dependent kinetochore phosphorylations promoting MPS1 kinetochore accumulation. Analysis of MPS1-GFP localization in cells briefly treated with Aurora B inhibitor (+AURKBi) confirmed that MPS1-GFP was lost from kinetochores upon Aurora B inhibition. This loss was not observed in cells that had been depleted of the PP2A catalytic α subunit or all PP2A-B56 regulatory subunits (Figure 2G and 2H). PP2A-B56 therefore counteracts the MPS1 kinetochore recruitment mediated by Aurora B. This effect appears to be dominant over the self-ejection triggered by the N-terminal phosphorylation of MPS1, explaining why total MPS1-GFP kinetochore accumulation is not reversed by depletion of PP2A-B56.

### MPS1-T675D/T676D is an active MPS1 kinase

To study the importance of MPS1 T-loop dephosphorylation by PP2A-B56 in isolation of the effect of PP2A-B56 on other substrates, we generated MPS1 mutants in which both the canonical T-loop residue T676 as well as the adjacent threonine T675 were mutated to phospho-mimetic aspartate (MPS1^DD^) or non-phosphorylatable alanine (MPS1^AA^). GFP-MPS1^AA^ and GFP-MPS1^DD^ mutant proteins were then analyzed for their ability to initiate and sustain spindle assembly checkpoint signaling alongside wild type (GFP-MPS1^WT^) and kinase-dead (GFP-MPS1^KD^) MPS1.

When endogenous MPS1 was replaced with the different forms of GFP-MPS1, GFP-MPS1^KD^ and GFP-MPS1^AA^ showed increased kinetochore levels in comparison to the wild type protein (Figure 3A and 3B, Figure 3G). This behavior of GFP-MPS1^KD^ and GFP-MPS1^AA^ is indicative of reduced or absent kinase activity (Hewitt et al., 2010; Jelluma et al., 2010). In contrast, GFP-MPS1^DD^ exhibited decreased levels of kinetochore recruitment (Figure 3A and 3B, Figure 3G). This observation is consistent with the idea that a constitutively active form of MPS1 should induce its own ejection from the kinetochore more effectively than the wild type protein (Wang et al., 2014). Only induction of GFP-MPS1^WT^ or GFP-MPS1^DD^ resulted in the effective recruitment of BUBR1 to the kinetochore and supported a cell cycle arrest in response to nocodazole (Figure 3A. 3C and 3D). All of these data taken together suggest that GFP-MPS1^DD^ represents an active form of MPS1.

**Figure 3.**
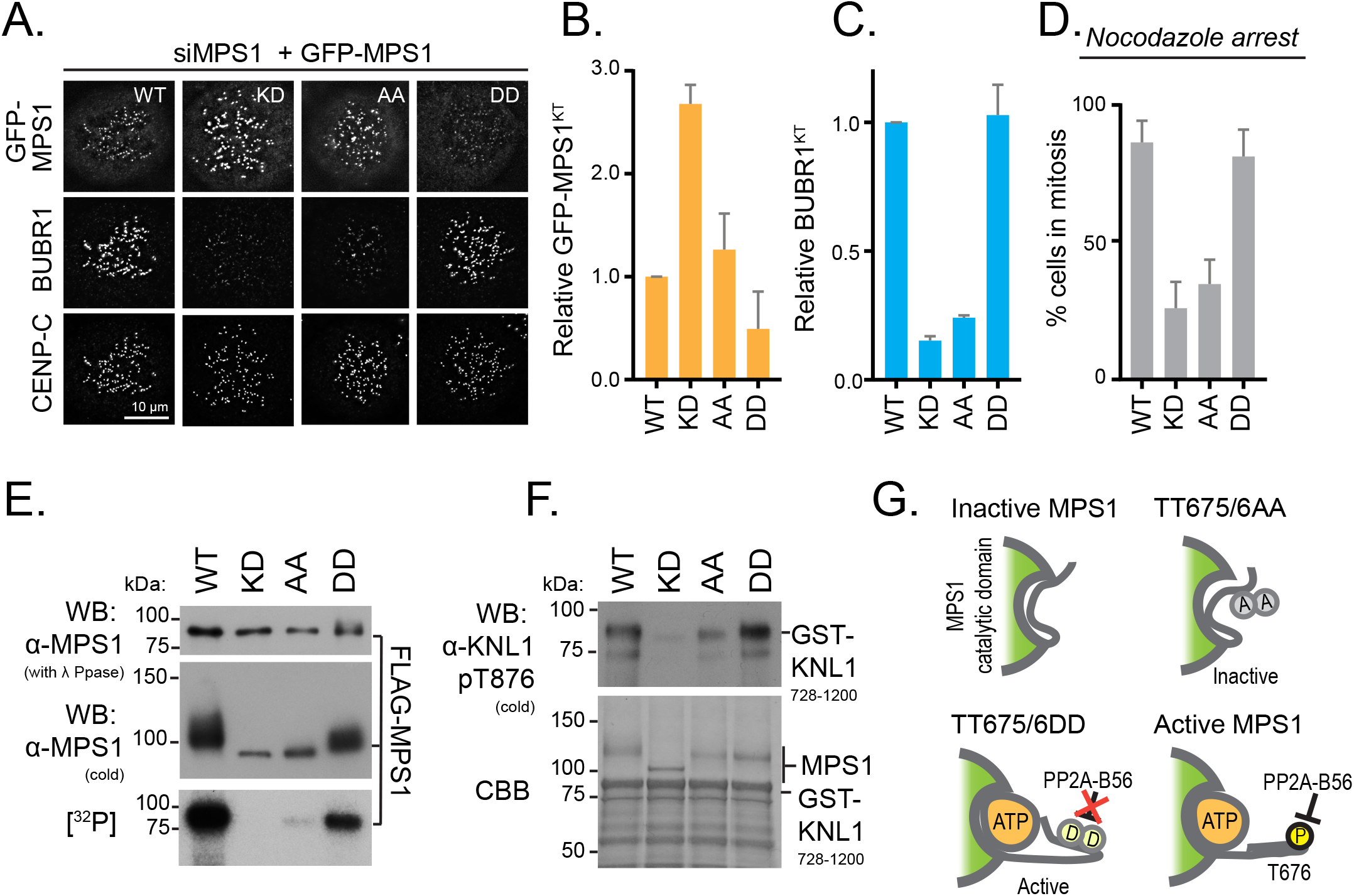
Replacing MPS1 T675 and T676 with Aspartic Acids restores MPS1 kinase activity and checkpoint functionality. **(A)** GFP-MPS1^WT^, MPS1^KD^, MPS1^AA^ or GFP-MPS1^DD^cells were arrested in mitosis. MPS1 was detected using GFP fluorescence, BUBR1 and CENP-C were detected using antibodies. Bar graphs show kinetochore associated **(B)** GFP-MPS and **(C)** BUBR1. Mean ± SEM for 20 kinetochores measured per cell in > 10 cells from 2 independent experiments. **(D)** The proportion of GFP-MPS1^WT^, MPS1^KD^, MPS1^AA^ or MPS1^DD^ expressing cells in mitosis after 16h 0.3 μM Nocodazole treatment is plotted in the bar graphs. Mean ± SEM for 2490 (WT), 2451 (KD), 2689 (AA) and 3211 (DD) from 2 independent experiments. **(E)** MPS1 autophosphorylation was measured by Western blotting and ^32^P incorporation. Equal loading of the different MPS1 forms was confirmed by Western blotting of λ-phosphatase treated proteins. **(F)** KNL1^728-1200^ phosphorylation by FLAG-MPS1 proteins was assessed by Western blotting with anti-KNL1^pT875^. Equal loading was confirmed by Coomassie Brilliant Blue (CBB) staining. **(G)** A cartoon illustrating how MPS1 TT675/6 mutations affect MPS1 activity.

Consistent with this proposal, purified FLAG-MPS1^WT^ and FLAG-MPS1^DD^ both exhibited a significant phosphorylation induced band upshift in Western blots which was not observed with kinase activity deficient FLAG-MPS1^KD^ and FLAG-MPS1^AA^ (Figure 3E). FLAG-MPS1^DD^ was less strongly upshifted than FLAG-MPS1^WT^, indicating that MPS1^DD^ has reduced kinase activity in comparison to wild type MPS1 (Figure 3E). Measurement of the autophosphorylation capacity of FLAG-MPS1^DD^ in comparison to FLAG-MPS1^WT^ in radioactive kinase assays indicated that FLAG-MPS1^DD^ possessed 42.60 ± 7.57 *%* of the kinase activity of wild type MPS1, in contrast to 0.72 ± 0.02 % for FLAG-MPS1^KD^ and 16.92 ± 6.01 % for FLAG-MPS1^AA^. FLAG-MPS1^DD^ phosphorylated a KNL1 fragment *in vitro* to near wild type levels (Figure 3F), and in all functional assays MPS1^DD^ kinase activity appeared to be sufficient to confer wild type levels of spindle assembly checkpoint proficiency (Figure 3A-3D, Figure 3G).

### Unregulated MPS1 activity traps cells in mitosis

To test the consequences of unregulated MPS1 activity for mitotic progression, HeLa-Flp-In/TREx cells depleted of endogenous MPS1 and expressing the different versions of GFP-MPS1 were filmed progressing through mitosis (Figure 4A and 4B). Replacement of endogenous MPS1 with GFP-MPS1^WT^ re-instated normal chromosome segregation and mitotic timing (Figure 4A, top panel; Figure 4C-4E). Expression of either GFP-MPS1^KD^ or GFP-MPS1^AA^ resulted in onset of chromosome segregation before completion of chromosome alignment and significantly shortened mitotic duration, consistent with a previous report (Jelluma et al., 2008a). These effects were more pronounced for GFP-MPS1^KD^ than for GFP-MPS1^AA^, in line with the idea that an MPS1-T-loop alanine mutation allows some residual kinase activity to take place (Figure 4A, second and third panel from top; Figure 4C-4E). Interestingly, cells expressing only the GFP-MPS1^DD^ version of MPS1 showed a phenotype distinct from both wild type as well as kinase dead and T-loop deficient MPS1. These cells entered mitosis normally but failed to align all their chromosomes and typically never reached a compact metaphase plate. The bulk of the chromosomes accumulated around a broad pseudo-metaphase with chromosomes continuously leaving this arrangement. Most of the cells remained trapped in this pseudo-metaphase state for several hours and eventually carried out an abnormal anaphase or apoptosed (Figure 4A, bottom panel, Figure 4C-4E).

**Figure 4.**
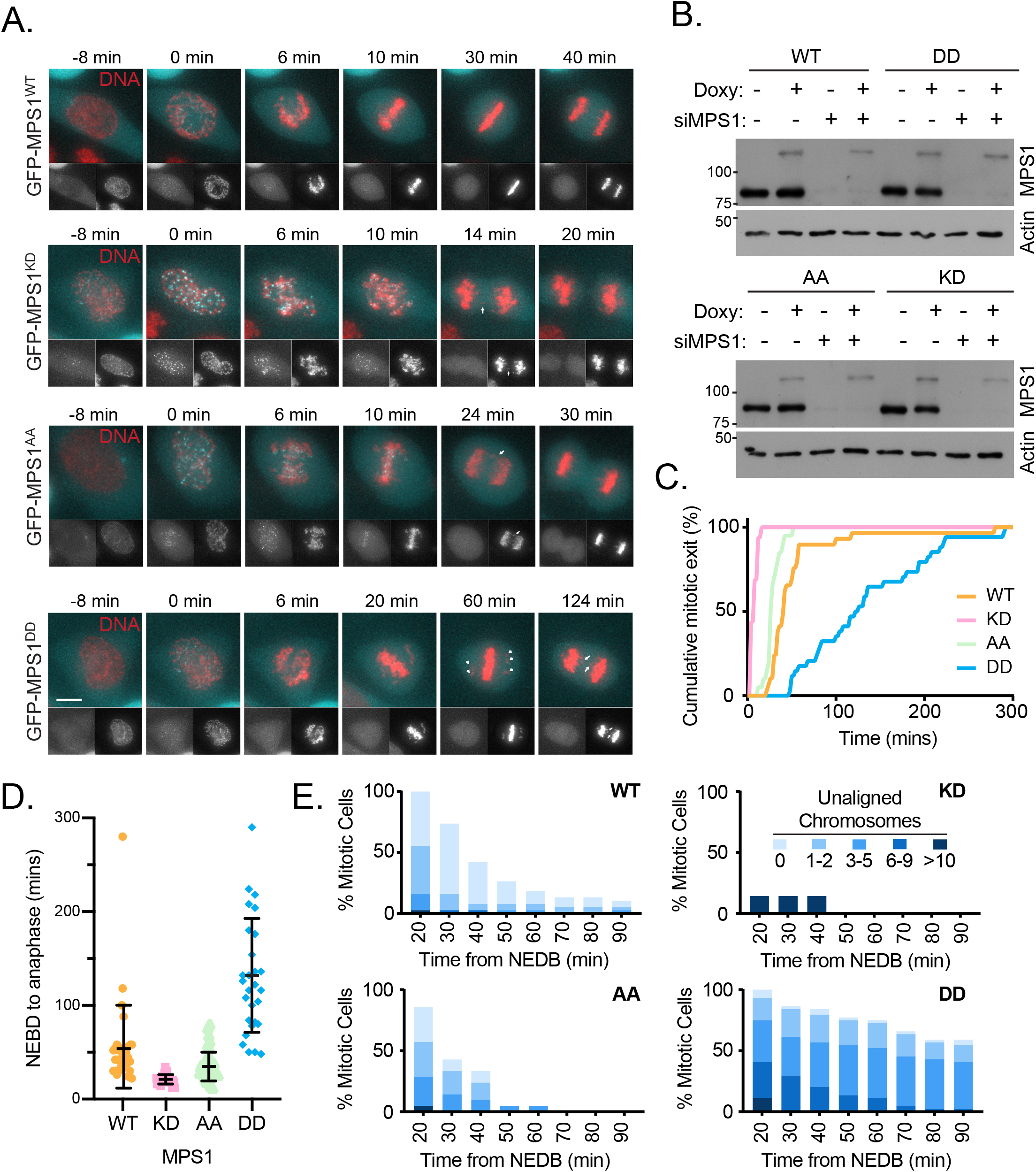
Expression of constitutively active MPS1 (TT675/6DD) results in extended mitotic duration. **(A)** Cells expressing GFP-MPS1 variants as indicated were imaged every 2 minutes as they passed through mitosis. GFP-MPS1 (Cyan and left hand insert) and DNA (red and right hand insert), are shown. Nuclear envelope breakdown is set to 0 min. Anaphase DNA bridges and unaligned chromosomes are indicated by arrows and arrowheads, respectively. **(B)** Cell lysates from cells in **(A)** were Western blotted for MPS1 and Actin (loading control). **(C)** Cumulative mitotic exit, **(D)** NEBD-anaphase duration and **(E)** the proportion of mitotic cells with unaligned chromosome were plotted over time. In **(D)** each point represents an individual cell with mean ± SD NEBD-anaphase time shown as bars. Data is from 2 independent experiments with 35 (WT), 27 (KD), 42 (AA) and 28 (DD) cells measured.

### Constitutively active MPS1 results in impaired microtubule-kinetochore attachment formation

In addition to its key role in orchestrating the spindle assembly checkpoint, MPS1 has been reported to be an important modulator of microtubule-kinetochore attachments (Jelluma et al., 2008b; Maciejowski et al., 2017; Santaguida et al., 2010). To understand the causes of the metaphase delay in cells with constitutively active MPS1, Hela-Flp-In/TREx cells depleted of endogenous MPS1 and expressing GFP-MPS1^WT^ or GFP-MPS1^DD^ were arrested in mitosis by MG132 treatment, and the alignment status of the chromosomes and checkpoint status of the kinetochores was evaluated. GFP-MPS1^DD^ cells exhibited significantly more unaligned chromosomes than GFP-MPS1^WT^ cells (Figure 5A and 5B) suggesting that microtubule-kinetochore attachments were compromised. Interestingly, only unattached kinetochores were decorated by GFP-MPS1^DD^ or MAD1 (Figure 5C), showing the behavior of the spindle checkpoint was not altered by the expression of the constitutively active form of MPS1 *per se*. To analyze chromosome alignment dynamics and error correction proficiency in more detail, a Monastrol washout assay was employed (Jelluma et al., 2008b). This analysis showed that unregulatable GFP-MPS1^DD^, like kinase inhibited MPS1, was defective in chromosome alignment upon Monastrol wash-out, with DD and AA and WT+MPS1i exhibiting significantly fewer aligned metaphase plates than WT (P < 0.05 (DD, AA), P < 0.005 (MPS1i) (Student’s *t* test)), and DD exhibiting significantly fewer aligned metaphase plates than AA (P < 0.05 (Student’s *t* test)) (Figure 5D-5F). The severity of the phenotype was more pronounced in the complete absence of MPS1 activity than in the unregulatable GFP-MPS1^DD^ mutant, possibly due to the different biological causes underlying the phenotype: in cells treated with MPS1 inhibitor, the main cause of chromosome alignment defects is the absence of MPS1-dependent PP2A-B56 recruitment to the kinetochore. Since this pool of PP2A-B56 opposes the error correction activities of both MPS1 and Aurora B, this results in unrestrained error correction. In contrast, cells expressing GFP-MPS1^DD^, are proficient in PP2A-B56 recruitment. However, because of the unchecked GFP-MPS1^DD^ kinase activity the phosphorylation-dephosphorylation balance of MPS1 targets in the error correction process is shifted towards phosphorylation and thus destabilization of attachments. Aurora B targets, however, are controlled normally in this situation.

**Figure 5.**
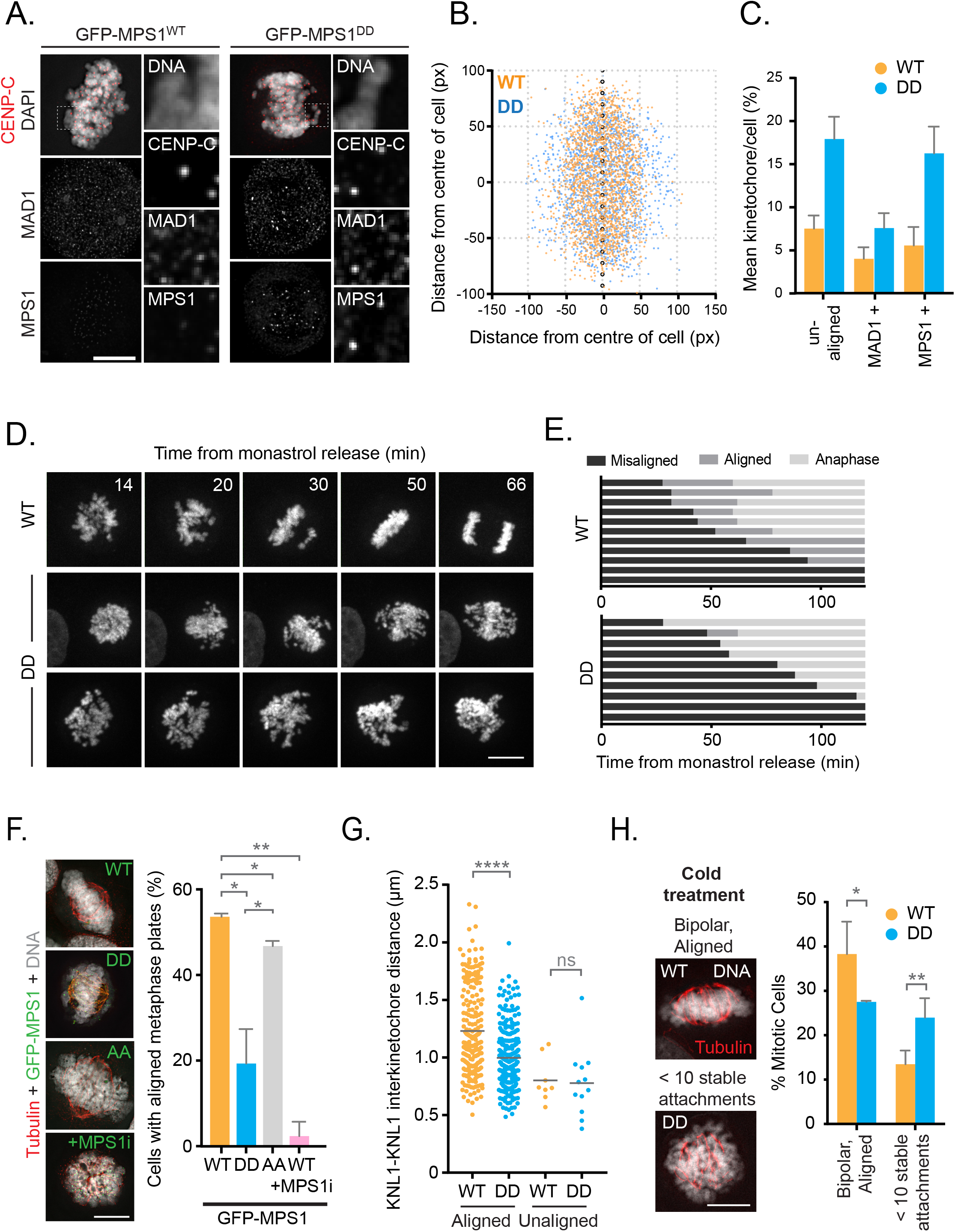
Microtubule-kinetochore interactions are perturbed in the presence of constitutively active MPS1. **(A)** MPS1^WT^ or GFP-MPS1^DD^ mitotic cells were stained for DNA (grey), MAD1 and CENP-C (red in the merged image). MPS1 was visualised by GFP fluorescence. An enlarged image of kinetochore pairs, indicated by a dashed box, is shown on the right-hand side of the images. **(B)** The x-y distance of individual kinetochores from the cell centre in 23 (MPS1^WT^) or 19 (MPS1^DD^) randomly selected cells is plotted. **(C)** Mean ± SEM proportion of misaligned, MAD1 positive or MPS1 positive kinetochores in the cells measured in **(B)**. **(D)** GFP-MPS1^WT^ or MPS1^DD^ expressing cells were filmed after Monastrol washout; images were captured every 2 min. Scale bar, 10 μm. **(E)** Quantification of cells from **(D)**. Each horizontal bar represents a single cell. **(F)** GFP-MPS1 expressing cells were fixed following a Monastrol washout into MG132. MPS1i was added at 2 μM for 60 min alongside MG132 where indicated. Images show DNA (grey), microtubules (red) and GFP-MPS1 (green). Bar graphs show mean proportion ± SD of cells exhibiting aligned metaphase plates. **(G)** KNL1-KNL1 inter-kinetochore distance is shown in GFP-MPS1^WT^ or GFP-MPS1^DD^ cells treated as in **(A)**. Each dot represents a kinetochore pair, and the black bar represents the mean average of 234 (WT) and 290 (DD) kinetochore pairs across 19 cells per condition from 2 independent experiments. **(H)** Cells as in **(A)** were cold treated prior to fixation. DNA (grey) and microtubules (red). The mean proportions ± SD of cells (282 WT and 331 DD cells from three independent experiments) containing bipolar spindles with aligned metaphase plates or cells with <10 stable microtubule-kinetochore attachments are plotted.

To demonstrate that the attachments that are formed in cells expressing GFP-MPS1^DD^ are less well stabilized than in control cells, the interkinetochore distances at aligned and unaligned chromosomes were measured in GFP-MPS1^WT^ and GFP-MPS1^DD^ cells. Indeed, the KNL1-KNL1 distance at aligned kinetochores in GFP-MPS1^DD^ cells was significantly smaller than (0.233 μm, p < 0.0005 (Student’s *t* test)) than in control cells, consistent with a reduction in pulling forces due to decreased microtubule-kinetochore attachment stability (Figure 5G).

To further test the idea that cells expressing GFP-MPS1^DD^ excessively turn over microtubule-kinetochore attachments, GFP-MPS1^DD^ and GFP-MPS1^WT^ cells were cold treated to selectively destabilize microtubules not attached to kinetochores. GFP-MPS1^DD^ cells exhibited significantly fewer aligned metaphase plates (P < 0.05 (Student’s *t* test)) and fewer cold-stable microtubules (P < 0.005 (Student’s *t* test)), confirming an impaired ability to stabilize microtubule-kinetochore attachments (Figure 5H). Taken together, the phenotype of the GFP-MPS1^DD^ expressing cells therefore seems to be primarily a consequence of uncontrolled MPS1-mediated error correction activity rather than unrestrained spindle assembly checkpoint activity (Jelluma et al., 2008b; Maciejowski et al., 2017).

MPS1 has two functions in the regulation of mitosis: it is the key regulator of the mitotic checkpoint, and at the same time it is also actively involved in error correction, a task which it shares with the Aurora B kinase (Pachis and Kops, 2018; Santaguida et al., 2010). We find here that unregulated kinase activity of MPS1 has a greater effect on error correction than on spindle assembly checkpoint control. One in-built safe-guarding mechanism that mitigates the consequences of hyperactive MPS1 is the fact that MPS1 activity is negatively correlated with its residence time at the kinetochore (Hewitt et al., 2010; Jelluma et al., 2010). A hyperactive MPS1 would therefore automatically be located less well to unattached kinetochores, the critical place for the initiation of spindle checkpoint signaling. Consistent with this idea we find that the phospho-mimetic GFP-MPS1^DD^ mutant shows significantly reduced kinetochore levels in comparison to wild type MPS1. The reduced MPS1 kinetochore residence time does not seem to affect spindle checkpoint signaling qualitatively, as cells expressing only GFP-MPS1^DD^ do not have any defects in initializing or maintaining a spindle checkpoint signal (Figure 3A-3D). This may suggest that unregulated MPS1 spindle checkpoint signaling for shorter time may amount to the same functional outcome as the wild type situation. However, interestingly, GFP-MPS1^DD^ cells do show signs of exaggerated error correction, apparent as the inability to align all chromosomes to a metaphase plate, reduced interkinetochore distances of aligned chromosomes and diminished numbers of cold-stable K-fibres (Figure 5). These observations are in line with the notion that MPS1 phosphorylates outer kinetochore targets, including the Ska complex to destabilize erroneous attachments (Maciejowski et al., 2017). One interesting idea is that, in contrast to the spindle checkpoint initiating phosphorylation of KNL1, these phosphorylation events may not be entirely dependent on the kinetochore localization of MPS1 and could be catalyzed by a cytoplasmic pool of MPS1. This would explain why for the regulation of kinetochore-microtubule attachments the unregulated nature of GFP-MPS1^DD^ is not offset by the shorter kinetochore residence time.

PP2A-B56 has been shown to counteract the microtubule-kinetochore attachment destabilizing activities of MPS1 as well as Aurora B (Foley et al., 2011; Kruse et al., 2013; Maciejowski et al., 2017; Suijkerbuijk et al., 2012; Xu et al., 2013). Using the same phosphatase to oppose both kinases thus allows coordinated stabilization of microtubule-kinetochore attachments. Our data indicate that PP2A-B56 not only dephosphorylates important targets of MPS1 in the error correction pathway but also regulates key regulatory residues on MPS1 itself. Intriguingly, in Drosophila, this latter function of dephosphorylating the MPS1 T-loop is carried out by PP1, not PP2A-B56, and seems to mainly affect MPS1’s role in controlling the spindle assembly checkpoint and not error correction, as Drosophila cells depleted of PP1 did not show any signs of elevated error correction (Moura et al., 2017). The reason for this difference to human cells may be found in the distinct wiring of some aspects of the spindle assembly checkpoint in flies in comparison to human cells. Most relevant for this discussion, it is not clear whether Drosophila has a distinct kinetochore pool of PP2A-B56. Certain functionalities of PP2A-B56 may therefore have been transferred to PP1 in flies. In human cells, PP2A-B56 emerges as the principal phosphatase opposing MPS1 phosphorylation events throughout mitosis, and our data further highlight the importance of this regulation.

## Materials and Methods

### Reagents and antibodies

General laboratory chemicals and reagents were obtained from Sigma-Aldrich and Thermo-Fisher Scientific unless specifically indicated. Inhibitors were obtained from Tocris Bioscience (MPS1-inhibitor AZ3146 20mM stock; PP1 and PP2A-inhibitors calyculin A 1mM stock, PP1-inhibitor tautomycetin, 2.5mM stock), Insight Bioscience (proteasome inhibitor MG132 20mM stock), Merck (microtubule polymerisation inhibitor nocodazole 6mM stock) and Cambridge Bioscience (Eg5 inhibitor Monastrol, 100mM stock). Inhibitor stocks were prepared in DMSO. Thymidine (Sigma Aldrich, 100mM stock) and doxycycline (Invivogen, 2mM stock) were dissolved in water. DNA vital dye SiR-Hoechst (Spirochrome) was dissolved in DMSO and used at 50 nM final concentration.

Commercially available polyclonal (pAb) or monoclonal (mAb) antibodies were used for MAD1 (Rabbit pAb; Genetex, GTX105079), BUBR1 (Rabbit pAb; Bethyl, A33-386A), beta Actin (HRP conjugated; Mouse mAb Abcam, [AC-15] ab49900), Tubulin (Mouse mAb; Sigma, [DM1A] T6199), PPP1CA (Rabbit pAb; Bethyl, A300-904A), PPP1CC (Goat pAb; Santa Cruz, sc6108), PP2CA(Goat pAb; Santa Cruz, SC6112), PP4C (Rabbit pAb; Bethyl, A300-835A), PP5C (Rabbit pAb; Bethyl, A300-909A), PP6C (rabbit pAb; Bethyl, A300-844A), PPP2R2A (mouse mAb; Cell Signaling, [2G9] 5689S), PPP2R5A (Rabbit pAb; Bethyl, A300-967A), PPP2R5D (Mouse mAb; Millipore, [H5D12] 04-639), PPP2R5E (Mouse mAb; Santa Cruz, [A-11] sc-376176), Phospho-MPS1T33/S37 (pRb; Thermo, 44-1325G), CENP-C (Guinea Pig pAb; MBL, PD030), FLAG epitope tag (Mouse mAb; ThermoFisher Scientific [FG4R], MA1-91878), GFP (Rabbit pAb; Abcam, ab290). Human CREST serum was obtained from Antibodies Inc (15-234-0001). Antibodies against MPS1 were raised in sheep (Scottish Blood Transfusion Services) against recombinant His-tagged MPS1 (amino acids 1-260) and affinity purified against the same recombinant protein. Antibodies to MPS1 phosphorylated at T676 were raised in sheep (Scottish Blood Transfusion Services) and affinity purified using the peptide sequence CMQPDTpTSVVKDS. Generation of anti-KNL1^pT875^ has been described previously (Espert et al., 2014). Secondary donkey antibodies against mouse, rabbit, guinea pig or sheep and labelled with Alexa Fluor 488, Alexa Fluor 555, Alexa Fluor 647, Cy5, or HRP were purchased from Molecular Probes and Jackson ImmunoResearch Laboratories, Inc., respectively. Affinity purified primary and HRP-coupled secondary antibodies were used at 1μg/ml final concentration. For western blotting, proteins were separated by SDS-PAGE and transferred to nitrocellulose using a Trans-blot Turbo system (Bio-Rad). Protein concentrations were measured by Bradford assay using Protein Assay Dye Reagent Concentrate (Bio-Rad). All western blots were revealed using ECL (GE Healthcare).

### Molecular biology

Human MPS1 was amplified from human testis cDNA (Marathon cDNA; Takara Bio Inc.) using Pfu polymerase (Agilent Technologies). MPS1 expression constructs were made using pcDNA5/FRT/TO vectors (Invitrogen) modified to encode the EGFP or FLAG reading frames. Mutagenesis was performed using the QuikChange method (Agilent Technologies). DNA primers were obtained from Invitrogen. Small interfering RNA (siRNA) duplexes targeting PPP family phosphatase subunits, BUBR1 and MPS1 have been described before (Espert et al., 2014; Hayward et al., 2019; Zeng et al., 2010). On-target SMARTPools were obtained from Dharmacon Horizon.

### Cell culture procedures

HeLa cells and HEK293T were cultured in DMEM with 1% [vol/vol] GlutaMAX (Life Technologies) containing 10% [vol/vol] bovine calf serum at 37°C and 5% CO_2_. For plasmid transfection and siRNA transfection, Mirus LT1 (Mirus Bio LLC) and Oligofectamine (Invitrogen), respectively, were used. HeLa cell lines with single integrated copies of the desired transgene were created using the T-Rex doxycycline-inducible Flp-In system (Invitrogen; (Tighe et al., 2004)).

CRISPR/Cas9-edited HeLa cells with an inserted GFP tag in the C-terminus of the TTK/MPS1 gene product and HeLa cells stably expressing GFP-MAD2 have been described before (Alfonso-Perez et al., 2019; Hayward et al., 2019).

### RNAi rescue assays

MPS1 siRNA rescue was performed by induction of GFP-MPS1 transgene for 6 h prior to a 48 h siRNA depletion of endogenous MPS1 using oligos against the 3’UTR (3’ UTR: 5’-UUGGACUGUUAUACUCUUGAA-3’, 5’-GUGGAUAGCAAGUAUAUUCUA-3’, and 5’-CUUGAAUCCCUGUGGAAAU-3’ (Hayward et al., 2019)). A second induction was performed 24 h into the siRNA depletion. GFP-BubR1 was induced 6 h prior to a 48h siRNA depletion of endogenous BubR1 using oligos against the 3’UTR (5’-GCAATCAAGTCTCACAGAT-3’ (Espert et al., 2014)). A second induction was performed 24 h into the siRNA depletion.

### Mitotic Arrests and Inhibitions

Unless otherwise stated, mitotically arrested cells were generated by addition of Nocodazole (0.3 μM, 2.5 h) followed by MG132 (20 μM, 0.5 h). Monastrol washouts (Figures 5D, 5E, 5F) were performed by adding Monastrol at 100 μM for 2.5 h, followed by a washing cells 3 times with warm PBS and 3 times with warm DMEM. In the case of Figure 5F, 20 μM MG132 was added for 60 min following washout. Cold treatment of cells (Figure 5H) was performed by incubating cells at 4 °C for 9 min prior to fixation to depolymerise microtubules that had not formed stable K-fibres. Inhibitor vehicle for all drugs was DMSO, and DMSO alone was added as control.

| **Inhibitor target** | **Inhibitor** | **Acronym used in text** | **Drug Concentration** | **Time of drug addition prior to fixation** |
| --- | --- | --- | --- | --- |
| MPS1 | AZ3145 | MPS1i | 2 μM | 5 min |
| PP1 | Tautomycetin | PP1i | 5 μM | 30 min |
| PP1 and PP2A | Calyculin A | PP1/PP2Ai | 25 nM | 6 min |
| Aurora B | ZM447439 | AURBK<sub>i</sub> | 2 μM | 10 min |

### Immunofluorescence microscopy and image processing

Cells were fixed with PTEMF (20mM PIPES-KOH pH 6.8, 0.2% [vol/vol] Triton X-100, 1mM MgCl2, 10mM EGTA, and 4% [wt/vol] formaldehyde)(Dunsch et al., 2011). Antibody dilutions were performed in PBS with 3% [wt/vol] BSA. Samples seeded on #1 thickness coverslips were imaged on a DeltaVision Core light microscopy system (GE Healthcare) using either a 60x/1.35na or 100x/1.4na objective fitted to an Olympus IX-71 microscope stand. Standard filter sets for DAPI (excitation 390/18, emission 435/48), FITC (ex. 475/28, em. 525/48), TRITC (ex. 542/27, em. 597/45) and Cy-5 (ex. 632/22, em. 676/34) were used to sequentially excite and collect fluorescence images on a CoolSnap HQ2 CCD camera (Photometrics) using the software package softWoRx (GE Healthcare). Cells were imaged using a 0.2μm interval and a total stack of 2μm and deconvolved for presentation using softWoRx. Image stacks were imported into FIJI (Schindelin et al., 2012) for maximum intensity projection and saving as 8-bit TIFF files. TIFF files were imported into Illustrator CS6 (Adobe) for figure production. For quantification imaging was performed using a 60x/1.35na oil immersion objective on a BX61 Olympus microscope equipped with filter sets for DAPI, EGFP/Alexa Fluor 488, Alexa Fluor 555, and Alexa Fluor 647 (Chroma Technology Corp.), a CoolSNAP HQ2 camera (Roper Scientific) and MetaMorph 7.5 imaging software (GE Healthcare).

### Quantification and Statistical Analysis

Image analysis was performed in FIJI and Excel (Microsoft). 10 Z-stacks with a 0.2 μm interval were sum projected for analysis. Relative protein kinetochore intensities (Figures 1B, 1C, 1E, 1F, 1H, 1I, 2B, 2C, 2E, 2F, 2H, 3B, 3C, S1D, S1E) were determined by placing a 10-pixel wide circular region of interest over individual kinetochores and measuring the mean pixel fluorescence, before dividing by the mean pixel intensity of the CENP-C channel within the same ROI. A mean background (cytoplasm) intensity for each cell is subtracted from each kinetochore protein or CENP-C measurement. The mean protein fluorescence of each kinetochore was divided by the mean kinetochore intensity of the total control population (always the condition closest to the Y-axis, with a value of 1) to generate relative values, which were plotted as bar graphs. > 15 cells per condition were analysed with ≥ 10 kinetochores per cell. All immunofluorescence experiments shown are representatives of at least two independent experiments pooled together. Unless otherwise stated, error bars represent the SEM, with N determined by the number of cells measured. Production of graphs was performed on GraphPad Prism (GraphPad Software, Inc.) using data exported from Excel. Statistical analysis of kinetochore intensities was carried out in Excel or GraphPad Prism.

### Inter-kinetochore distance

KNL1-KNL1 inter-kinetochore distance measurements were taken in a threedimensional space from 234 (WT) and 290 (DD) kinetochore pairs across 19 cells per condition from 2 independent experiments. Distances between a kinetochore pair are plotted as individual points, with the mean distance plotted as a black bar. Students T-test was performed to determine statistical significance.

### Kinetochore positioning measurements

Kinetochore positions were identified as the centre of mass of CENP-C signals identified within the cell boundary following equal thresholding of images, and were plotted in Figure 5B by distance in pixels from the centre of the cell. 23 (WT) or 19 (DD) randomly selected cells were measured. There was no significant difference (Student’s T-test) between the number of identified kinetochores per cell in WT (mean=91, S.D.=17) and DD (mean=94, S.D.=15). The alignment status of these kinetochores (Figure 5C) was determined as defining aligned kinetochores as being positioned within the 50% region of the X-axis closest to the centre of a cell (i.e. within 25% either side of the centre of the cell along the X-axis). Misaligned kinetochores are therefore defined as being outside of this zone. Proportion of MAD1 and MPS1 positive kinetochores were quantified as number of objects within the cell boundary counted following equal thresholding of image channels.

### Live cell microscopy and statistical analysis

Time-lapse imaging of cells with a paired control sample was performed on a DeltaVision Elite light microscopy system as described for fixed cell samples. Fluorescence images were collected on a 512×512 pixel EMCCD camera (QuantEM, Photometrics) using the software package softWoRx (GE Healthcare). Cells were placed in a 37°C and 5% CO_2_ environmental chamber (Tokai Hit) on the microscope stage with lens heating collar. Cells were seeded on 2-chambered glass bottomed dishes (Lab-Tek) at 30,000 per well. SiR-Hoechst (Spirochrome) was added 8h prior to imaging at a final concentration of 100nM. Typically, 7 planes were captured per cell 2μm apart every 2min with laser powers at 2% and 25ms exposures. Deconvolution and maximum intensity projections were performed using softWoRx, with image cropping performed using FIJI.

All other time-lapse imaging was performed using an Ultraview Vox spinning disc confocal system (Perkin Elmer) mounted on an Olympus IX81 inverted microscope, a 512×512 pixel EMCCD camera (ImagEM C9100-13, Hamamatsu Photonics) and Volocity software. Cells were placed in a 37°C and 5% CO2environmental chamber (Tokai Hit) on the microscope stage with lens heating collar. Imaging was performed using a 60x NA1.4 oil immersion objective, 4−12% laser power and 30-200ms exposure time. Typically, 19 planes, 0.6μm apart were imaged every 2min. Maximum intensity projection or summed projection of the fluorescent channels was performed in FIJI. Statistical analysis of live cell imaging data (Figure 2C, Figure 4D) was carried out in GraphPad Prism.

### Protein expression and purification

FLAG-MPS1^WT^, FLAG-MPS1^KD^, FLAG-MPS1^AA^ and FLAG-MPS1^DD^ were expressed and purified from HEK293T cells. For each construct, two 15 cm dishes of cells were transfected with 8μg DNA, each, for 36h, including a 12h nocodazole arrest. Cell pellets were lysed in 1ml lysis buffer (20mM Tris-HCl pH 7.4, 300mM NaCl, 1% [vol/vol] Triton X-100 and protease inhibitor cocktail (Sigma-Aldrich)). MPS1 was immunoprecipitated from the clarified supernatants using 100μl FLAG-agarose beads (Sigma-Aldrich). IPs were washed twice with lysis buffer, four times with 20mM Tris-HCl pH 7.4, 300mM NaCl, 0.1% [vol/vol] Triton X-100, two times with 20mM Tris-HCl pH 7.4, 300mM NaCl and once with 100mM Tris-HCl pH 7.4. GST-tagged KNL1^728-1200^ was expressed and purified as described before (Espert et al., 2014).

### MPS1 kinase assays

For kinase assays, 1μg recombinant GST-Knl1^728-1200^ was phosphorylated with 1μg recombinant FLAG-MPS1 on FLAG-agarose beads for 30min at 30°C in 50mM Tris-HCl pH 7.3, 50mM KCl, 10mM MgCl_2_, 20mM sodium β-glycerophosphate, 15mM EGTA, 0.2mM ATP (cold assay) or 0.1mM ATP (hot Assay), 1mM DTT, and 1μCi [^32^P]γ-ATP (hot assay) per reaction.

Incorporation of γ^32^P into MPS1 or pKNL1^875^ intensity was used as a readout of kinase activity. Coomassie stained bands of FLAG-MPS1 with incorporated γ^32^P on an SDS-PAGE were excised for scintillation counting. The mean proportion ± SD of MPS1 construct activity (cpm) relative to WT was determined across 2 independent experiments. The values were then normalised to the number of available phosphorylation sites to account for the fact that FLAG-MPS1^AA^ and FLAG-MPS1^DD^ were lacking two sites. These two proteins were considered to have ten available sites and FLAG-MPS1^WT^ and FLAG-MPS1^KD^ twelve (Dou et al., 2011; Tyler et al., 2009).

### Online supplemental material

Figure S1 shows the specificity of the MPS1 pT676 antibody, the full phosphatase catalytic subunit RNAi screen for pT676 retention upon MPS1 inhibition and Western blots demonstrating efficient phosphatase and BUBR1 depletion as well as efficient BUBR1 transgene induction, supporting Figure 1. Figure S2 shows pT676 staining in control, PP1 or PP2A-B56 depleted cells with intact microtubules, supporting Figure 1. Videos 1-4 show cells expressing the different GFP-MPS1 mutants progressing through the cell cycle, supplementing Figure 4.

## Supporting information

Supplemental video GFP-MPS1-AA

Supplemental video GFP-MPS1-DD

Supplemental video GFP-MPS1-KD

Supplemental video GFP-MPS1-WT

## Acknowledgements

DH was supported by a Medical Research Council Senior Non-Clinical Research fellowship awarded to UG (MR/K006703/1) and TAP by a Biotechnology and Biological Sciences Research Council Strategic LoLa grant (BB/M00354X/1). We thank Stephen Taylor (University of Manchester) for Hela-Flp-In/TREx cells, Jakob Nilsson (University of Copenhagen) for helpful advice regarding phospho-mimetic mutants and Iona Manley and Zoë Geraghty for critical reading of the manuscript.

The authors declare no competing financial interests.

## Author Contributions

Conceptualization: U. Gruneberg. Investigation: D. Hayward, J. Bancroft, T. Alfonso-Pérez, D. Mangat, S. Dugdale and J. McCarthy. Funding acquisition: U. Gruneberg, F. A. Barr. Supervision: U. Gruneberg, F. A. Barr. Writing – original draft: U. Gruneberg. Writing − review and editing: T. Alfonso-Pérez, D. Hayward, J. Bancroft, F. A. Barr and U. Gruneberg.

