## Supplemental material for "Checkpoint signaling and error correction require regulation of the MPS1 T-loop by PP2A-B56"

### Supplemental Figure legends

**Figure S1. MPS1 autophosphorylation and localisation are regulated by PP2A-B56. (A)** Control depleted or MPS1-depleted HeLa MPS1-GFP mitotic cells were treated with MPS1 inhibitor or DMSO for 5 min as indicated. Cells were stained with antibodies against MPS1 pT676 and CENP-C. Total MPS1-GFP was detected by GFP fluorescence. (**B**) Recombinant, insect cell expressed wild type (WT) or kinase-dead (KD) His-MPS1 was treated with λ-phosphatase or reaction buffer alone and analysed by SDS-PAGE and Coomassie Brilliant Blue staining (CBB) or Western blotting with antibodies against MPS1 pT676. **(C)** MPS1 T-loop phosphorylation and endogenous MPS1 levels were assessed in control depleted mitotic HeLa MPS1-GFP cells or HeLa MPS1-GFP cells depleted of PP1αβγ (siPP1), PP2CA (siPP2A), PPP4C (siPP4), PPP5C (siPP5) or PPP6C (siPP6) and treated with MPS1i as indicated. Mean kinetochore MPS1 pT676 phosphorylation **(D)** and MPS1-GFP kinetochore levels ± SEM **(E)** relative to CENP-C are plotted. Values are from three independent experiments with 15 kinetochores measured in at least 10 cells per condition. **(F)** Representative Western blot analysis of PP1, PP2A, PP4, PP5 and PP6 depletion efficiencies. Actin is shown as a loading control. **(G)** Representative Western blot of PP2A-B55 and PP2A-B56 depletion efficiencies. Actin is used as a loading control. **(H)** Western blot of HeLa-Flp-In/TREx GFP-BUBR1$^{WT}$ or GFP-

BUBR1$^{L669A/I672A}$ cells either with (+) or without (-) BUBR1 3'-UTR siRNA and/or transgene induction (Doxy), blotted for anti-BUBR1 and anti-GFP. Actin is shown as a loading control.

**Figure S2. PP1 depletion in an unperturbed cell cycle does not lead to retention of MPS1 T-loop phosphorylation**

(**A**) HeLa cells stably expressing GFP-MAD2 were control depleted or depleted of PP$\alpha\gamma$ (siPP1) or all PP2A-B56 subunits for 48 h and fixed and stained with antibodies against MPS1 pT676. Kinetochores were detected with CREST serum and MAD2 by GFP fluorescence. A spindle assembly checkpoint positive kinetochore in siPP1 depleted cells is indicated by a small arrow. Scale bar, 10 µm. **(B)** MPS1 pT676 levels ± SEM relative to CENP-C were quantitated and plotted. Values are from two independent experiments with 15 kinetochores measured in at least 10 cells per condition.

**Videos 1-4**

HeLa cells, depleted of endogenous MPS1 and expressing GFP-MPS1 variants (cyan) were filmed progressing through mitosis. SiR-Hoechst was added 8 h before imaging to visualise DNA. Image stacks were captured every 2 mins. Nuclear envelope breakdown is marked at 0 mins. Videos are shown at 7 fps.

**Figure S1** Hayward et al.

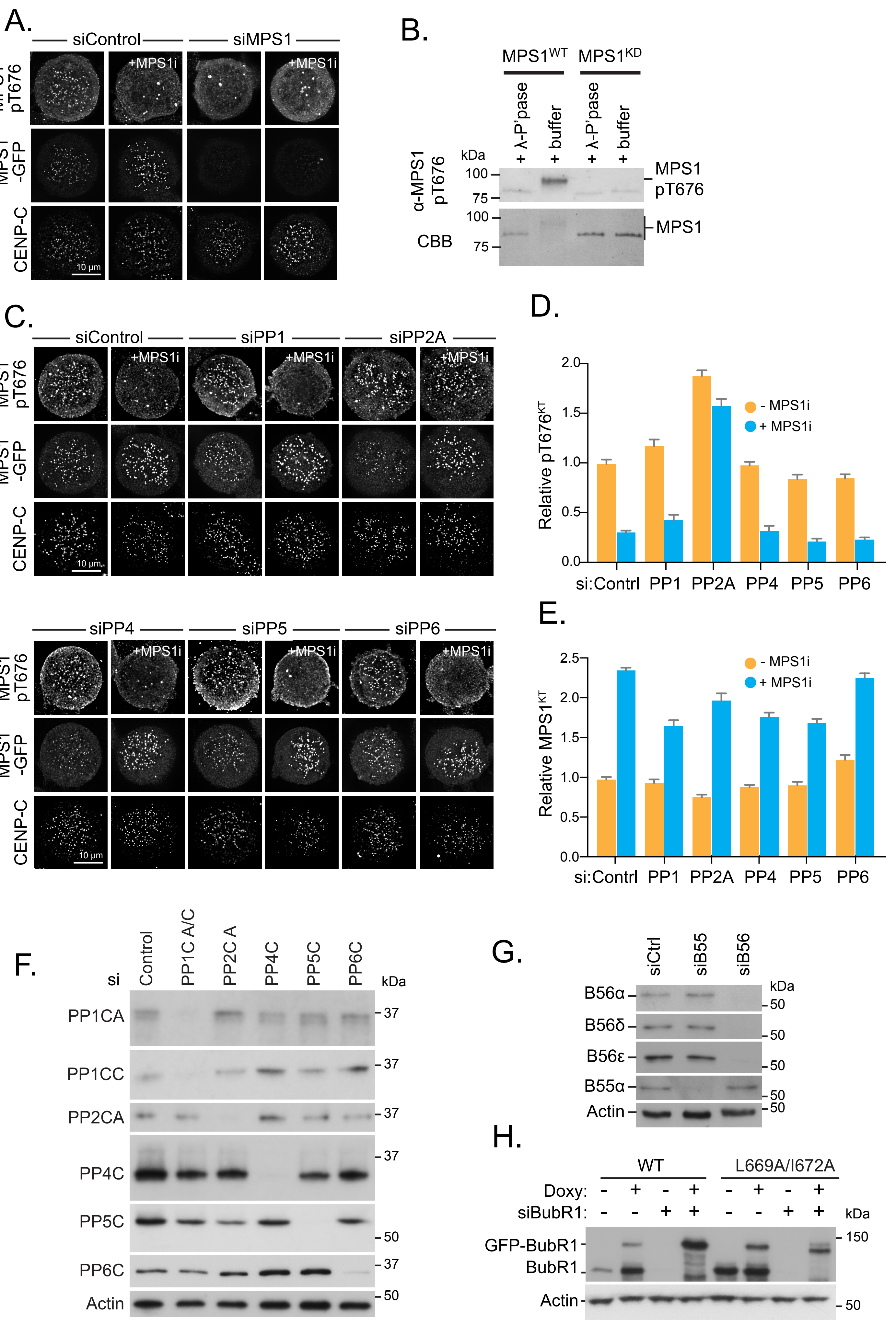

**Figure S2** Hayward et al.

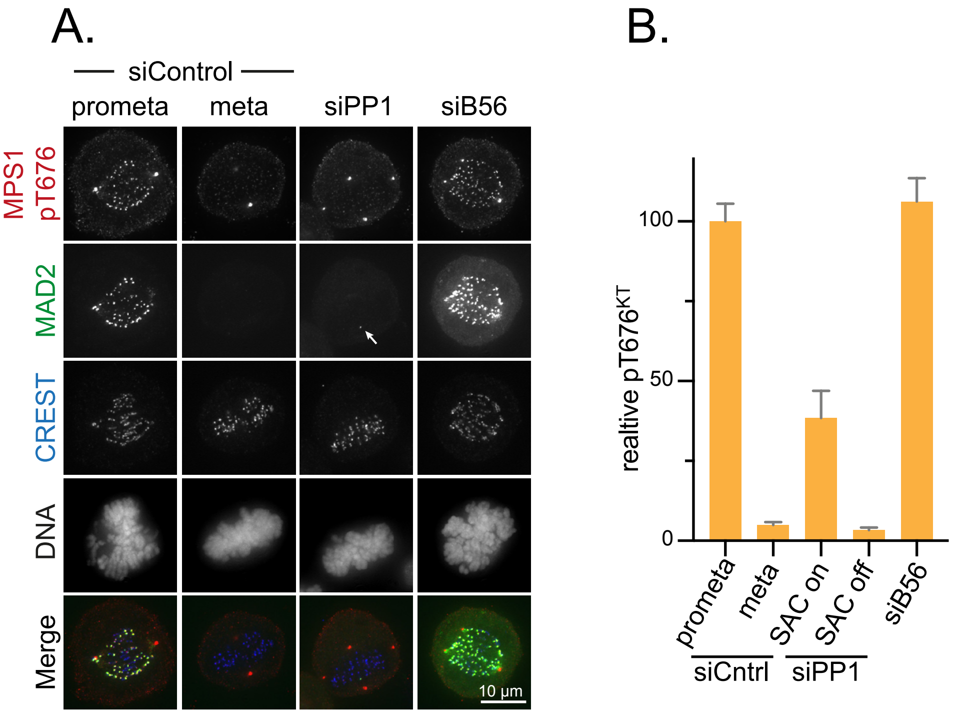
